# Nucleolar translocation of human DNA topoisomerase II by ATP depletion and its disruption by the RNA polymerase I inhibitor BMH-21

**DOI:** 10.1101/2021.09.29.462339

**Authors:** Keiko Morotomi-Yano, Ken-ichi Yano

## Abstract

DNA topoisomerase II (Top2) is a nuclear protein that resolves DNA topological problems and plays critical roles in multiple nuclear processes. Human cells have two Top2 proteins, Top2A and Top2B, that are localized in both the nucleoplasm and nucleolus. Previously, ATP depletion was shown to augment the nucleolar localization of Top2B, but the molecular details of subnuclear distributions, particularly of Top2A, remained to be fully elucidated in relation to the status of cellular ATP. Here, we analyzed the nuclear dynamics of human Top2A and Top2B in ATP-depleted cells. Both proteins rapidly translocated from the nucleoplasm to the nucleolus in response to ATP depletion. FRAP analysis demonstrated that they were highly mobile in the nucleoplasm and nucleolus. The nucleolar retention of both proteins was sensitive to the RNA polymerase I inhibitor BMH-21, and the Top2 proteins in the nucleolus were immediately dispersed into the nucleoplasm by BMH-21. Under ATP-depleted conditions, the Top2 poison etoposide was less effective, indicating the therapeutic relevance of Top2 subnuclear distributions. These results give novel insights into the subnuclear dynamics of Top2 in relation to cellular ATP levels and also provide discussions about its possible mechanisms and biological significance.

## Introduction

DNA topoisomerase II (Top2) is an enzyme that controls the topological state of DNA ^1,2^. Top2 plays multiple roles in the nucleus, because DNA topological constraints arise from various nuclear processes ^3,4^. Human and mammalian cells have two Top2 proteins, Top2A and Top2B, which are encoded by distinct genes ^5,6^. Both proteins consist of an N-terminal ATPase domain, a central catalytic domain, and a C-terminal region ^2^. Top2A and Top2B share high sequence similarity in their ATPase and catalytic domains and exhibit similar enzymological properties *in vitro* ^7,8^. In a cell, Top2A and Top2B are differentially expressed and participate in different nuclear processes. Top2A is mainly expressed in the S and G2 phases of the cell cycle and thus exists in proliferating cells ^9,10^. Top2A plays critical roles in DNA replication and chromosome segregation, and the genetic knockout of Top2A leads to lethality at a cellular level ^4,11^. Top2B is expressed throughout the cell cycle and is present in both dividing and non-dividing cells ^10^. Top2B plays roles in the transcription of hormone-inducible genes ^12-14^ and repair of DNA double strand breaks ^15^. Although Top2B is dispensable for cell proliferation, its physiological importance is evident in non-dividing cells, where Top2A is absent. Top2B is involved in the control of neuronally expressed genes ^16^, and its genetic knockout causes defects in neural development, resulting in neonatal lethality ^17^.

Top2A and Top2B are mobile in the nucleus and are localized in both the nucleoplasm and nucleolus ^18-21^. Domain analyses of Top2A and Top2B identified a short region that is responsible for the nucleolar localization of each protein ^20,22^. These regions are located between the catalytic domain and the C-terminal region and act as RNA-binding sites ^20,22^.

Intriguingly, the subnuclear distribution of Top2B is affected by the energy status of the cell. Pharmacological inhibition of ATP synthesis leads to the accumulation of Top2B in the nucleolus ^20^. Similarly, membrane permeabilization using mild detergent, which causes extracellular leakage of ATP, results in augmented localization of Top2B in the nucleolus ^15,23^. These observations collectively indicate that Top2B is a prominent example of an ATP-dependent shift in subnuclear protein distribution.

The nucleolus is a major site of ribosomal biogenesis and is assembled around tandem repeats of ribosomal DNA ^24^. The nucleolus contains highly mobile components, many of which can shuttle between the nucleoplasm and nucleolus, because of the membrane-less structure and liquid-like nature of the nucleolus ^25^. In addition to ribosomal biogenesis, the nucleolus participates in other cellular functions, including cell cycle control, stress response, and DNA damage response ^26^. Accordingly, the nucleolus contains many proteins that exert their primary functions in the nucleoplasm and appear to be unrelated to ribosomal biogenesis ^27^. The subnuclear distribution of these proteins is often affected by pharmacological and physical treatment of cells. For example, the proteasome inhibitor MG132 induces the translocation of p53 and MDM2 from the nucleoplasm to the nucleolus, and this MG132-induced nucleolar translocation is suppressed by PI3-kinase inhibitors, such as wortmannin and LY-294002 ^28^. Extracellular acidosis results in the translocation of VHL from the nucleoplasm to the nucleoli ^29^. Heat shock induces the increased nucleolar localization of the Polycomb group protein CBX ^30^. Conversely, PARP1, WRN, XRCC1, and TARG1, all of which are involved in DNA repair, are translocated from the nucleolus to the nucleoplasm in response to DNA damage induction and oxidative stress ^31-34^. The subnuclear distribution of WRN is controlled by the enzymatic activities of ATM, HDAC, and Sirt1, and the pharmacological inhibition of these factors perturbs the proper subnuclear localization of WRN ^35,36^.

A growing number of studies suggest that transient shifts in the equilibrium of protein distribution between the nucleoplasm and nucleolus may be a widespread phenomenon and may serve to protect against cellular insults ^37-39^. As for Top2, the ATP dependency of Top2B subnuclear distribution was previously described ^20^, but the underlying molecular details, particularly those for Top2A, remain to be fully understood in relation to the cellular ATP status. In this study, we sought to explore how human Top2A and Top2B respond to ATP depletion. Our observations shed further light on Top2 as an example where cellular ATP levels profoundly affect subnuclear protein distribution.

## Results

### ATP depletion induces the translocation of Top2A and Top2B to the nucleoli

Although Top2B was reported to accumulate in the nucleoli of ATP-depleted cells, the precise relationship between ATP levels and the subnuclear distribution of Top2 remains to be clarified. Thus, we initiated our experiments by measuring ATP levels. Simultaneous inhibition of glycolytic and mitochondrial ATP-producing reactions is known to lead to rapid ATP depletion ^40^. Thus, we treated HeLa cells with 2-deoxy-glucose (2DG) and carbonyl cyanide m-chlorophenylhydrazone (CCCP), which are a hexokinase inhibitor and a mitochondrial oxidative phosphorylation uncoupler, respectively ^41,42^. As shown in Fig. 1a, we observed a rapid decrease in intracellular ATP levels within 5 min after 2DG/CCCP treatment.

**Fig. 1.**
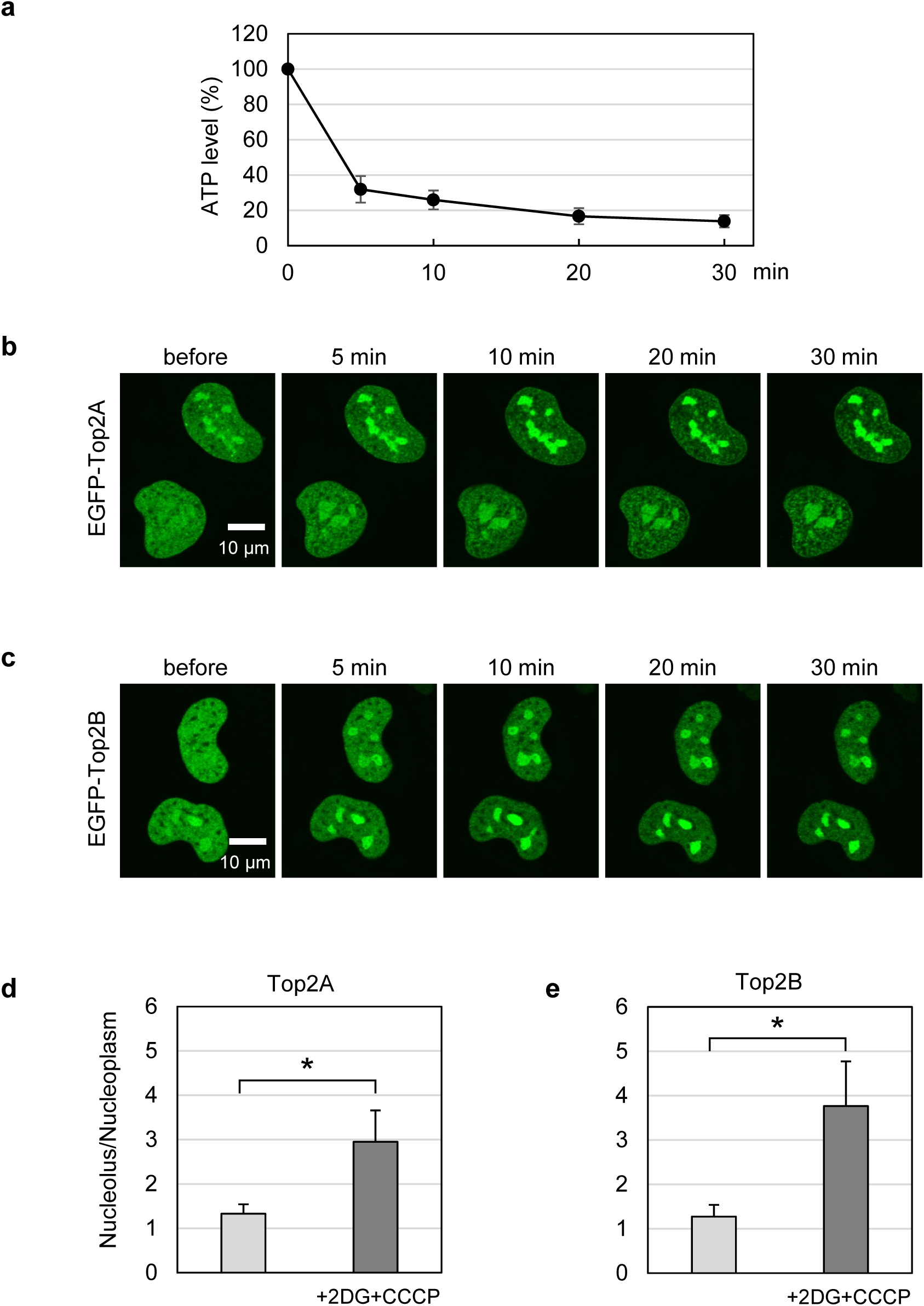
ATP depletion induces translocation of EGFP-Top2A and EGFP-Top2B to the nucleoli. (**a**) Intracellular ATP levels after 2DG/CCCP treatment. HeLa cells were treated with 25 mM 2-deoxyglucose and 10 μM CCCP, and intracellular ATP levels were measured at the indicated time points. Average values with SD from 6 independent experiments were plotted. (**b**) Translocation of EGFP-Top2A to the nucleoli by ATP depletion. Cells expressing EGFP-Top2A were treated with 25 mM 2DG and 10 μM CCCP, and images were captured at the indicated time points. (**c**) Translocation of EGFP-Top2B to the nucleoli by ATP depletion. An experiment was performed on EGFP-Top2B as described in (**b**). (**d, e**) Nucleolus-to-nucleoplasm ratios of EGFP fluorescence. Cells expressing EGFP-Top2A (**d**) or EGFP-Top2B (**e**) were incubated in the presence and absence of 25 mM 2DG and 10 μM CCCP, and images were captured at 20 min. Green fluorescence in the nucleolus and nucleoplasm was quantified, and average values of the nucleolus-to-nucleoplasm ratio and SD were calculated (n=30; *: statistically significant, **d**: p=7.97E-14, **e**: p=1.23E-14).

Next, we analyzed the effects of 2DG/CCCP treatment on the subnuclear distribution of EGFP-tagged Top2 proteins in HeLa cells. Before 2DG/CCCP treatment, EGFP-Top2A (Fig. 1b) and EGFP-Top2B (Fig. 1c) were localized in both the nucleoplasm and nucleoli. Within 5 min of 2DG/CCCP addition, we observed increased fluorescence signals in the nucleoli and reduced nucleoplasmic signals of both EGFP-Top2 proteins, which became increasingly evident over time (Fig 1b, c). These observations indicate that 2DG/CCCP treatment induced the translocation of EGFP-Top2 proteins from the nucleoplasm to the nucleoli. We calculated the nucleolus-to-nucleoplasm ratios of fluorescence before and after 2DG/CCCP treatment and quantitatively validated the translocation of EGFP-Top2 proteins after 2DG/CCCP treatment (Fig. 1d, e).

Next, we carried out immunofluorescence staining for endogenous Top2 proteins along with the nucleolar protein fibrillarin (Fig. 2). Top2A is expressed in a cell cycle-dependent manner ^9^, and Top2B is constitutively expressed throughout the cell cycle ^10^. Accordingly, the intensity of Top2A staining differed from cell to cell, presumably reflecting the cell cycle status of individual cells (Fig. 2a). Top2B staining was relatively uniform, owing to the constitutive expression of Top2B (Fig. 2b). Without 2DG/CCCP treatment, both the nucleoplasm and nucleoli were immunostained for Top2A and Top2B. Cells treated with 2DG/CCCP exhibited stronger staining in the nucleoli than in the nucleoplasm (Fig. 2a, b). These results suggest that endogenous Top2A and Top2B translocate from the nucleoplasm to the nucleoli in ATP-depleted cells, which is consistent with the observations of EGFP-Top2A and EGFP-Top2B in living cells (Fig. 1).

**Fig. 2.**
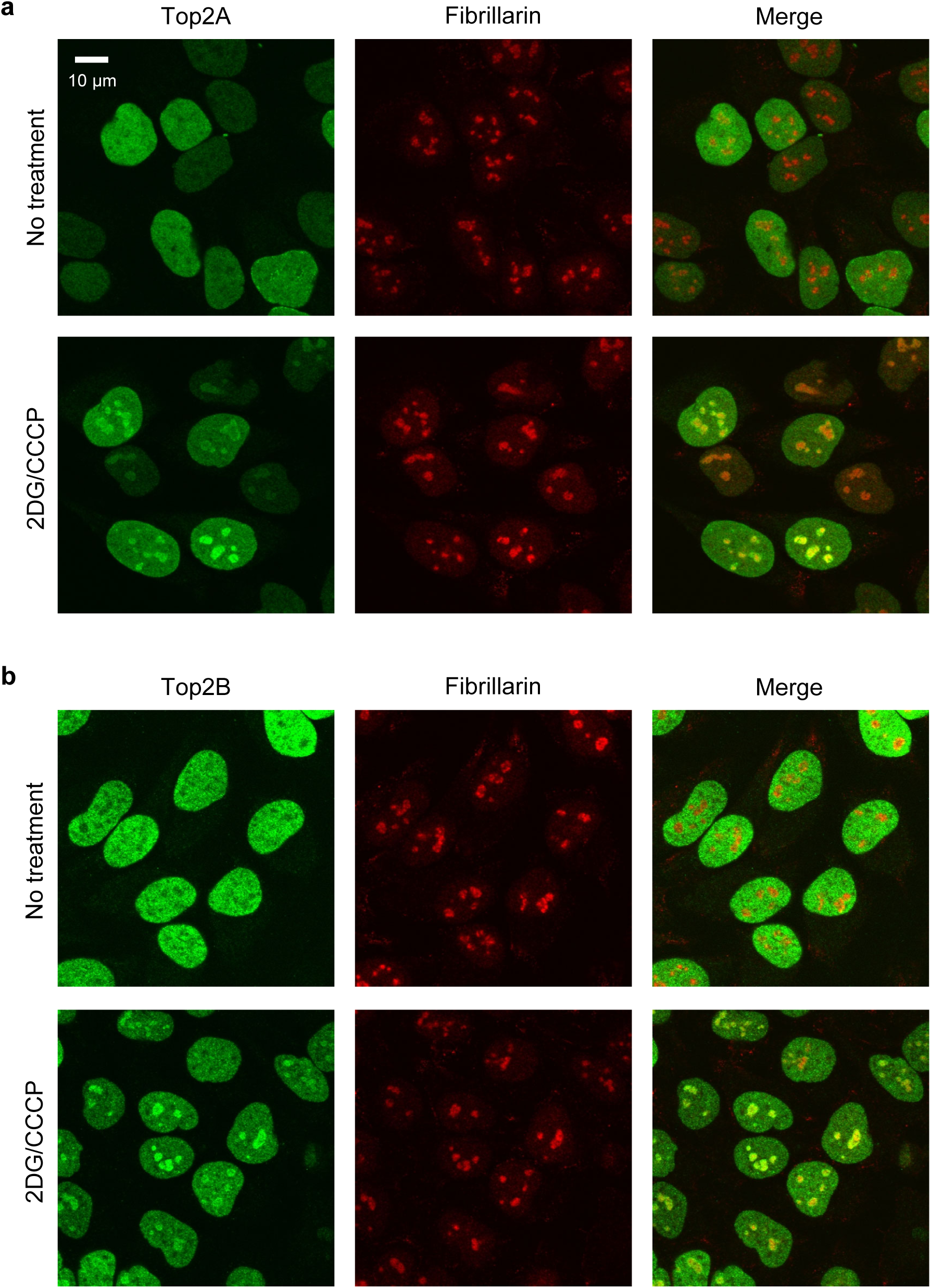
Endogenous Top2A and Top2B proteins are localized to the nucleoli in ATP-depleted cells. (**a**) Immunofluorescence staining for endogenous Top2A and fibrillarin. HeLa cells were incubated with or without 25 mM 2DG/10 μM CCCP for 30 min. Cells were subjected to coimmunostaining for endogenous Top2A (green) and fibrillarin (red). (**b**) Coimmunostaining for endogenous Top2B (green) and fibrillarin (red) was performed as described in (**a**).

### EGFP-Top2A and -Top2B are highly mobile in the nucleoplasm and nucleoli

We next performed FRAP analysis to explore how Top2 proteins exist in the nucleoplasm and nucleolus. Without 2DG/CCCP treatment, a small area in the nucleoplasm of an EGFP-Top2A-expressing cell was photobleached (Fig 3a). We observed rapid recovery of fluorescence in the bleached nucleoplasm, implying that EGFP-Top2A is highly mobile in the nucleoplasm. Next, using a 2DG/CCCP-treated cell, we carried out a photobleaching experiment in the nucleolus and observed rapid recovery of fluorescence after photobleaching, suggesting the high mobility of EGFP-Top2A in the nucleolus of ATP-depleted cells (Fig. 3b). Quantification of fluorescence signals before and after photobleaching further demonstrates that EGFP-Top2A is highly mobile in both the nucleoplasm (Fig. 3c) and nucleolus (Fig. 3d).

**Fig. 3.**
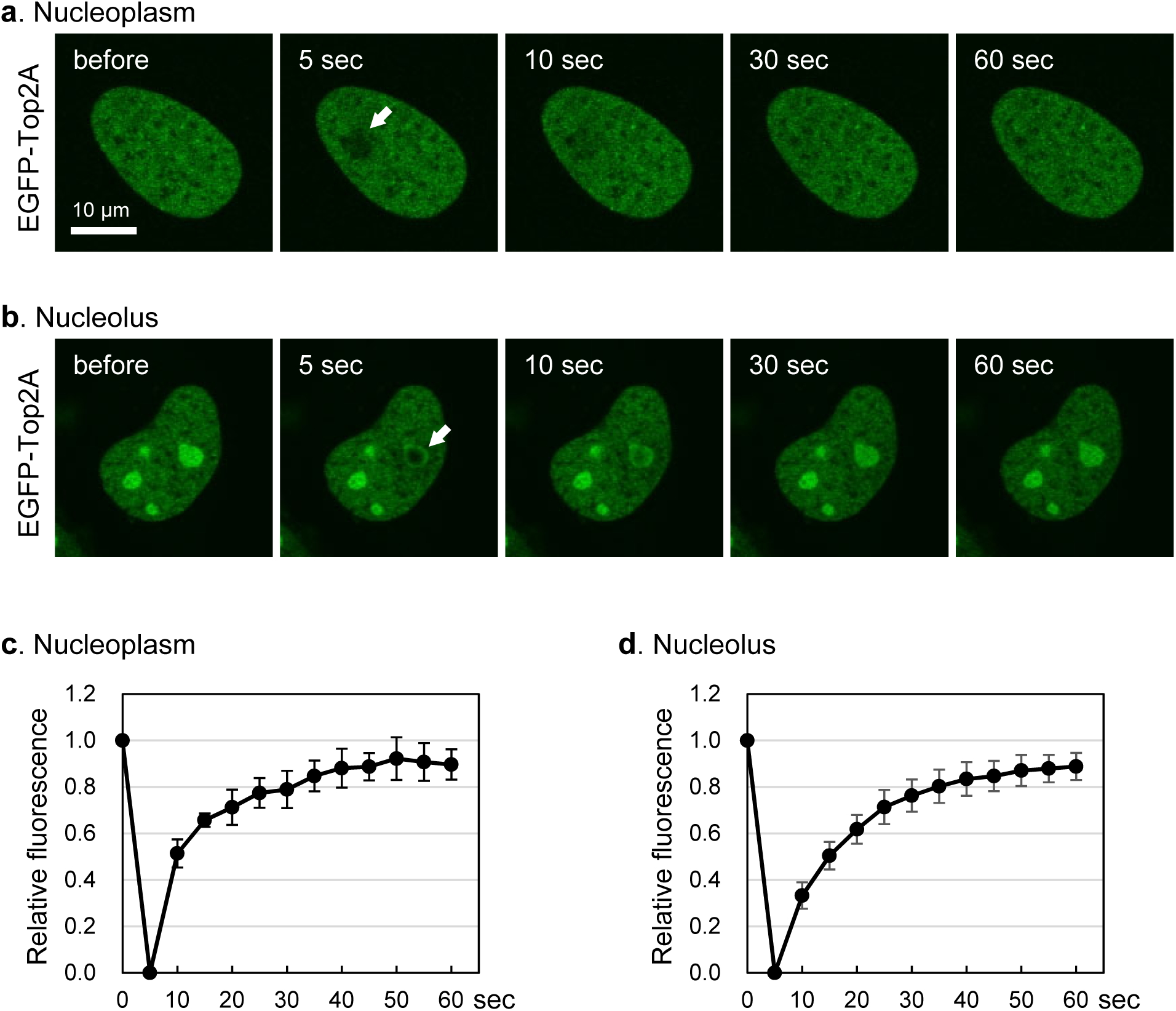
EGFP-Top2A is highly mobile in the nucleoplasm and nucleolus. (**a**) Representative images of FRAP analysis of EGFP-Top2A in the nucleoplasm. A small area of the nucleoplasm (shown with a white arrow) of a HeLa cell expressing EGFP-Top2A was photobleached, and fluorescence images were captured at the indicated time points. (**b**) Representative images of FRAP analysis of EGFP-Top2A in the nucleolus. A HeLa cell expressing EGFP-Top2A was incubated with 25 mM 2DG and 10 μM CCCP for 20 min, and photobleaching in a small area of the nucleolus (shown with a white arrow) was carried out. (**c, d**) Quantification of fluorescence signals of EGFP-Top2A in the nucleoplasm (**c**) and in the nucleolus (**d**). Fluorescence images were captured at 5 sec intervals. Photobleaching was conducted at 5 sec. Average values of relative fluorescence and SD were calculated (n=10).

We performed similar analysis on EGFP-Top2B and observed rapid recovery of fluorescence in the nucleoplasm (Fig 4a) and nucleolus (Fig 4b) after photobleaching. We confirmed the high mobility of Top2B in the nucleoplasm (Fig. 4c) and nucleoli (Fig. 4d) by fluorescence quantification.

**Fig. 4.**
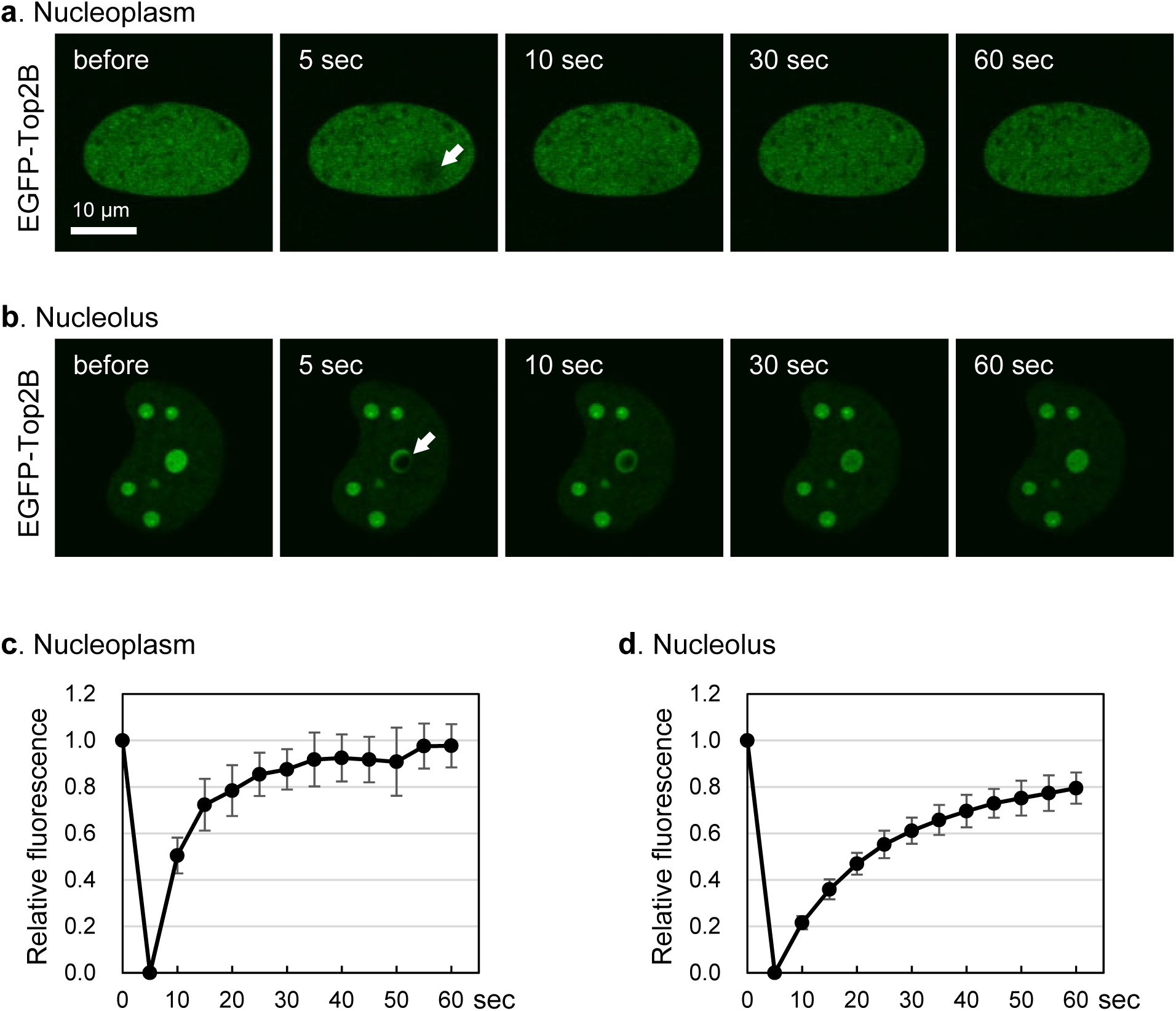
EGFP-Top2B is highly mobile in the nucleoplasm and nucleolus. Experiments were performed on EGFP-Top2B as described in Fig. 3. (**a, b**) Representative images of FRAP analysis of EGFP-Top2B in the nucleoplasm (**a**) and in the nucleolus (**b**). (**c, d**) Quantification of fluorescence signals of EGFP-Top2B in the nucleoplasm (**c**) and in the nucleolus (**d**).

### Pretreatment with the RNA polymerase I inhibitor BMH-21 suppresses the nucleolar translocation of Top2A and Top2B in ATP-depleted cells

To gain insights into the mechanism underlying Top2 nucleolar translocation, we attempted pharmacological inhibition of the nucleolar translocation of Top2 using various compounds.

Although most tested compounds had no obvious effects on the Top2 nuclear distribution with or without ATP depletion, we found that pretreatment with the RNA polymerase I inhibitor BMX-21 suppressed the nucleolar translocation of Top2 in ATP-depleted cells. First, we compared the effects of inhibitors of three RNA polymerases (Pol) on EGFP-Top2A (Fig. 5a) and EGFP-Top2B (Fig. 5b). Cells were pretreated with one of the Pol inhibitors and subsequently treated with 2DG/CCCP in the presence of respective inhibitors. Pretreatment with BMH-21 (Pol I inhibitor) completely suppressed the nucleolar translocation of Top2 by 2DG/CCCP addition. In contrast, α-amanitin (Pol II inhibitor) and ML-60218 (Pol III inhibitor) did not affect the nucleolar translocation of EGFP-Top2 in 2DG/CCCP-treated cells. Quantification of fluorescence signals in the nucleolus and nucleoplasm verified the suppression of Top2 nucleolar translocation by BMH-21 pretreatment (Fig. 5c, d). We performed immunostaining of endogenous Top2 proteins and fibrillarin and further confirmed that BMH-21 pretreatment suppressed the nucleolar translocation of Top2 (Fig. 5e, f).

**Fig. 5.**
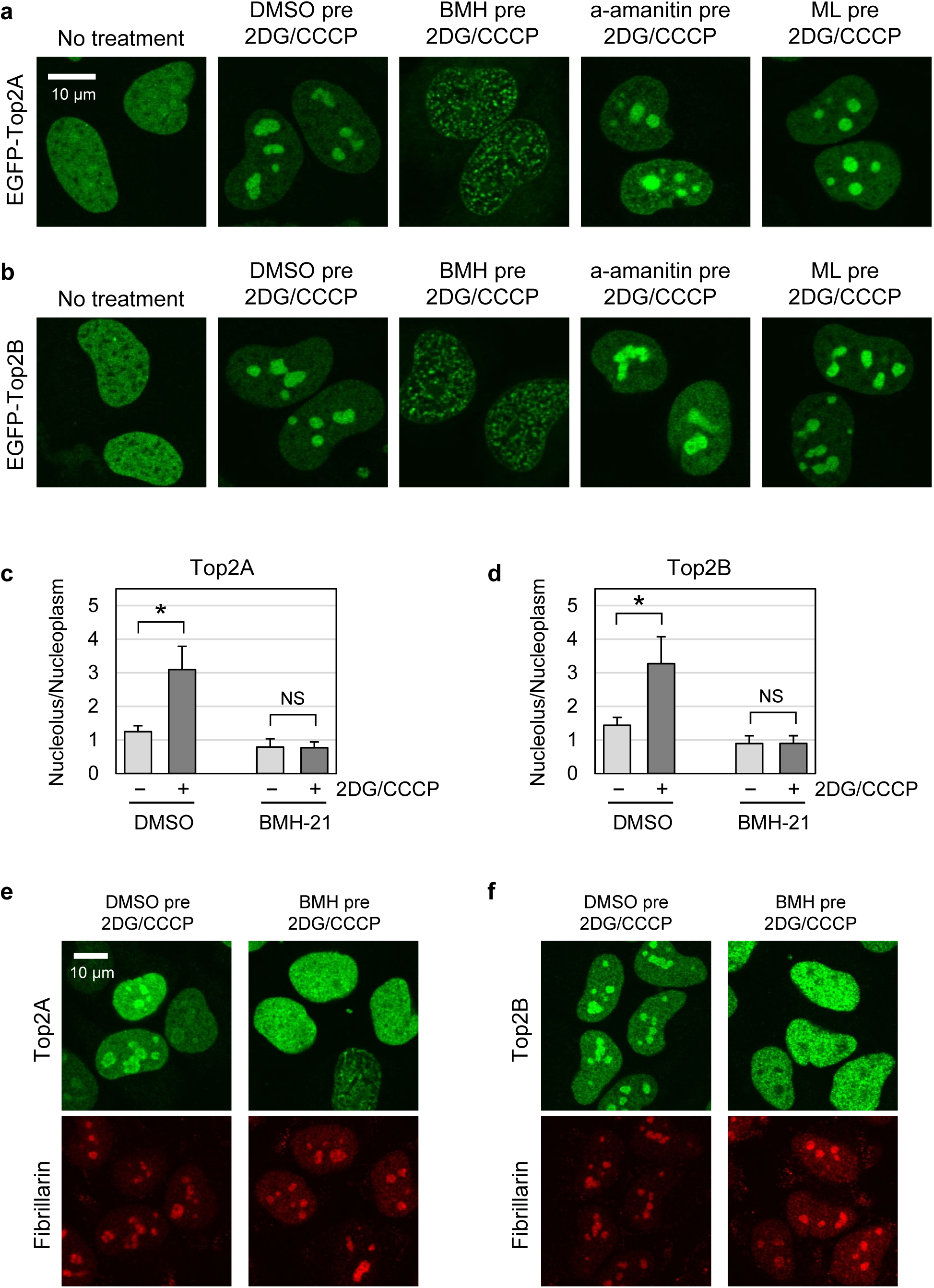
The RNA polymerase I inhibitor BMH-21 suppresses the nucleolar translocation of Top2A and Top2B in ATP-depleted cells. (**a, b**) Effects of RNA polymerase inhibitors on the nucleolar translocation of EGFP-Top2A (**a**) and EGFP-Top2B (**b**) in ATP-depleted cells. HeLa cells expressing EGFP-Top2A or EGFP-Top2B were pretreated with either DMSO (vehicle, 30 min), BMH-21 (1 μM, 30 min), α-amanitin (10 μg/ml, 120 min), or ML-60218 (50 μM, 60 min). Cells were subsequently treated with 25 mM 2DG and 10 μM CCCP in the presence of respective RNA polymerase inhibitors for 15 min. (**c**) Nucleolus-to-nucleoplasm ratios of EGFP-Top2A. Cells expressing EGFP-Top2A were pretreated with either DMSO or 1 μM BMH-21 and subsequently treated with 2DG/CCCP as described in (**a**). Green fluorescence in the nucleolus and nucleoplasm was quantified, and average values of the nucleolus-to-nucleoplasm ratios and SD were calculated. (n=30; *: statistically significant, p=1.31E-15; NS: Not significant, p=0.714) (**d**) Nucleolus-to-nucleoplasm ratios of EGFP-Top2B. Experiments were performed on EGFP-Top2B as described in (**c**). (n=30; *: statistically significant, p=7.82E-14; NS: Not significant, p=0.973) (**e, f**) Immunofluorescence staining for endogenous Top2A (**e**) and Top2B (**f**). HeLa cells were treated as described in (**a**) and subjected to immunostaining for either Top2A or Top2B. Fibrillarin was coimmunostained to indicate the nucleoli (red).

### BMH-21 rapidly abolishes the nucleolar accumulation of EGFP-Top2 and -Top2B in ATP-depleted cells

As shown in Fig. 1, EGFP-Top2A and EGFP-Top2B were stably retained in the nucleoli during 2DG/CCCP treatment. Next, we investigated the effect of BMH-21 on the nucleolar retention of EGFP-Top2A. Cells were treated with 2DG/CCCP to allow the translocation of EGFP-Top2A from the nucleoplasm to the nucleoli, yielding nucleolar accumulation of EGFP-Top2A. The cells were then treated with BMH-21 in the presence of 2DG/CCCP (Fig. 6a). As shown in Fig. 6b, the accumulation of EGFP-Top2A in the nucleoli was rapidly abolished by the addition of 1 μM BMH-21. Within 1 min after BMH-21 addition, EGFP-Top2A in the nucleoli was significantly decreased. At 2 min, EGFP-Top2A evenly distributed in the nucleoplasm and nucleoli (Fig. 6b). Furthermore, when the concentration of BMH-21 was increased from 1 μM to 2 μM, we observed faster dispersion of EGFP-Top2A from the nucleoli to the nucleoplasm (Fig. 6c). We confirmed no effects of vehicle (DMSO) treatment in the same experimental procedure (Fig. 6d). Quantification of the nucleolus-to-nucleoplasm ratios validated the rapid disruption of nucleolar accumulation of EGFP-Top2A by BMH-21 (Fig. 6e, f).

**Fig. 6.**
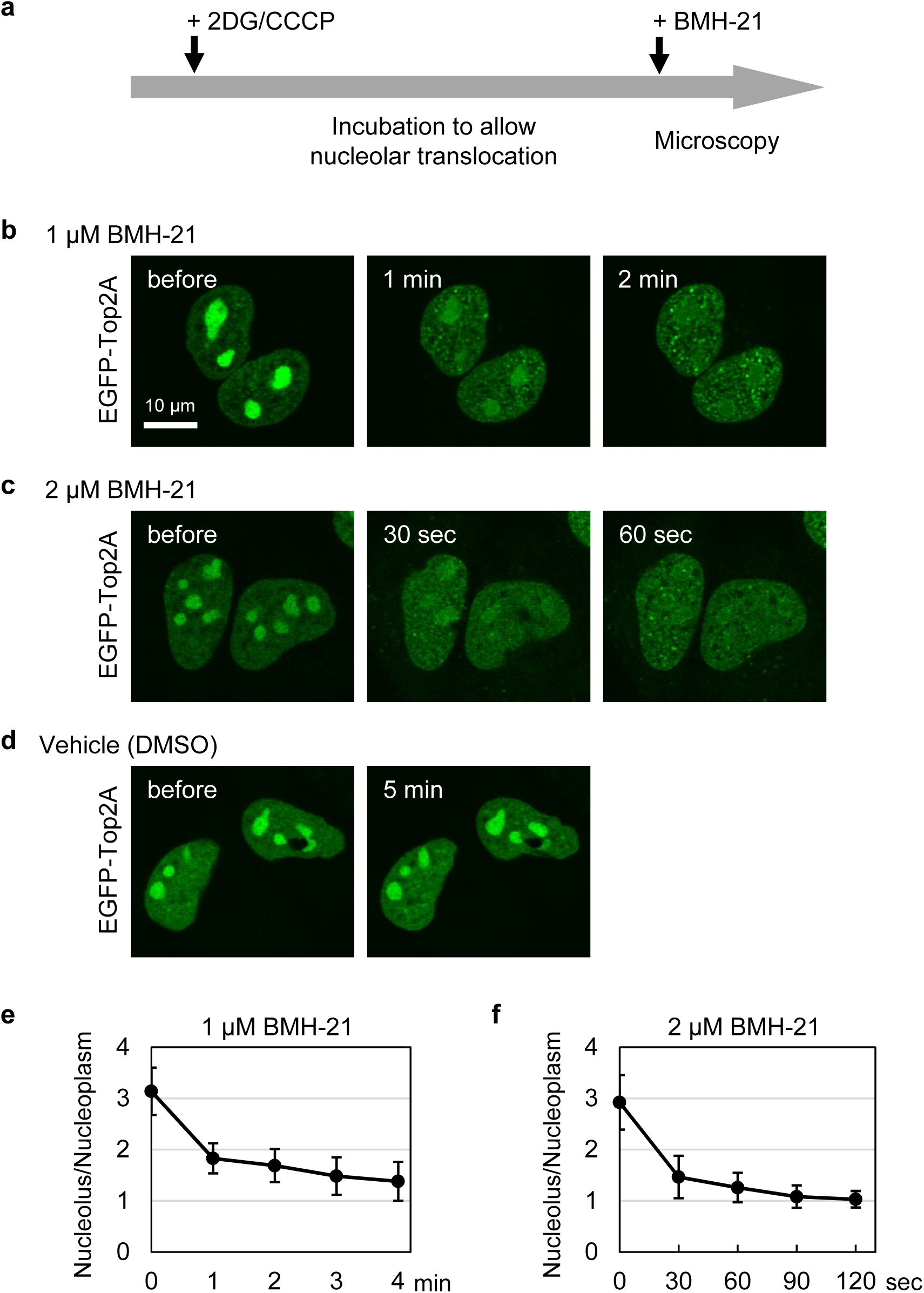
BMH-21 disrupts the retention of EGFP-Top2A in the nucleolus. (**a**) An experimental scheme for drug addition and fluorescence microscopy. (**b, c**) Rapid dispersion of nucleolar EGFP-Top2A by BMH-21. HeLa cells expressing EGFP-Top2A were incubated with 25 mM 2DG/10 μM CCCP for 20 min to allow translocation of EGFP-Top2A to the nucleolus. Cells were subsequently treated with either BMH-21 at 1 μM (**b**) or 2 μM (**c**) in the presence of 2DG/CCCP. Fluorescence images were captured at the indicated periods. (**d**) A control experiment using DMSO, instead of BMH-21. (**e, f**) Quantification of the nucleolus-to-nucleoplasm ratios of EGFP-Top2A. Experiments were performed using 1 μM (**e**) or 2 μM (**f**) BMH-21 as described in (**b**) and (**c**). Average values and SD were calculated (n=10).

We repeated the experiments on EGFP-Top2B and observed that BMH-21 rapidly abolished the nucleolar accumulation of EGFP-Top2B (Fig. 7a, b, c), which was quantitatively verified as shown in Fig. 7d and 7e. Of note, two major nucleolar proteins, nucleolin and fibrillarin, were consistently localized in the nucleoli during treatment with 2DG, CCCP, and BMH-21 (Fig. 7f, g).

**Fig. 7.**
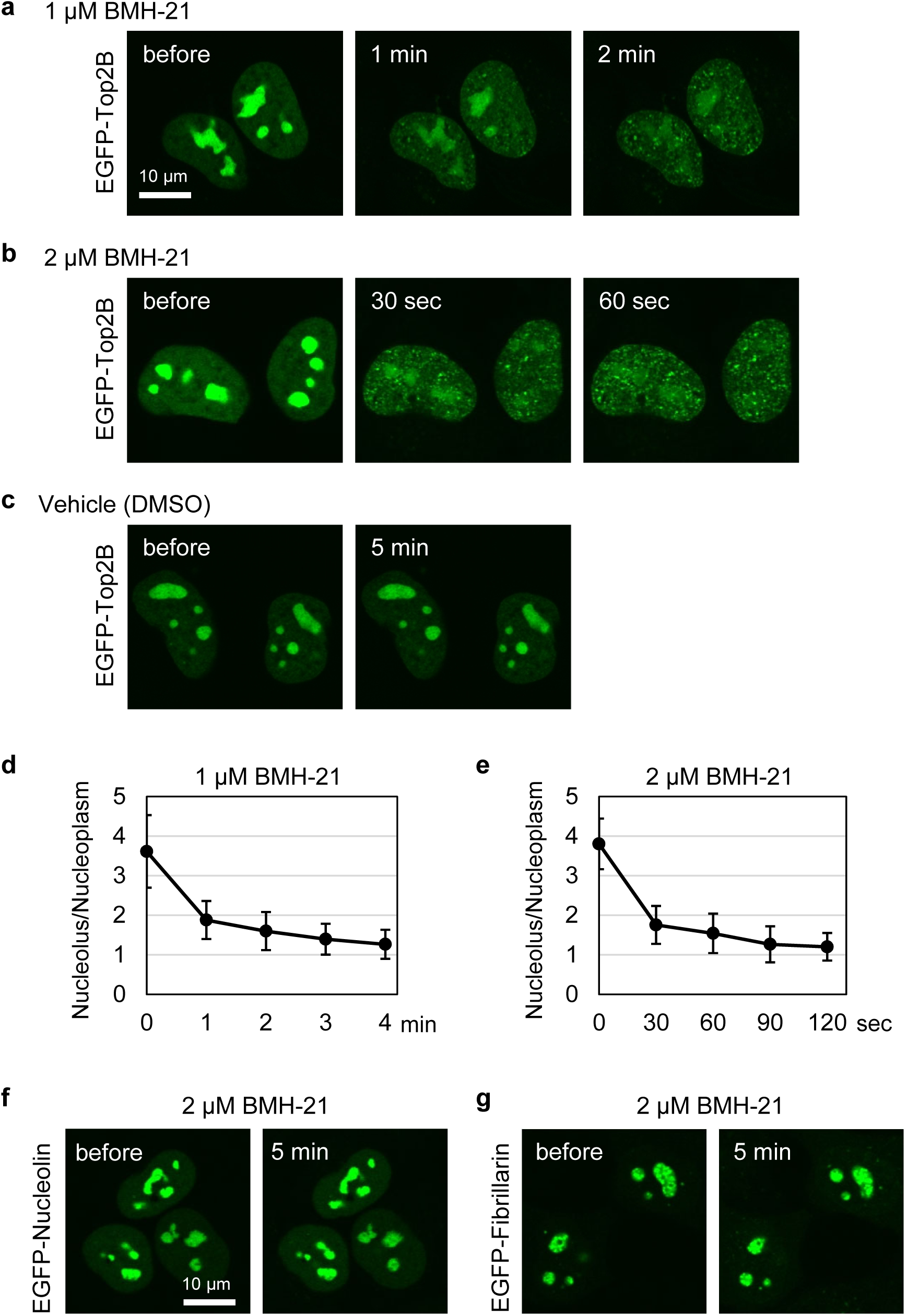
BMH-21 disrupts the retention of EGFP-Top2B in the nucleolus. (**a, b**) Rapid dispersion of nucleolar EGFP-Top2B by BMH-21. Experiments were performed on EGFP-Top2B as described in Fig. 6 b and c. (**c**) A control experiment using DMSO, instead of BMH-21. (**d, e**) Quantification of the nucleolus-to-nucleoplasm ratios of EGFP-Top2B. Experiments were performed using 1 μM (**d**) or 2 μM (**e**) BMH-21. Average values and SD were calculated (n=10). (**f, g**) BMH-21 treatment does not affect the nucleolar localization of nucleolin and fibrillarin in ATP-depleted cells. HeLa cells expressing either EGFP-nucleolin (**f**) or EGFP-fibrillarin (**g**) were incubated with 25 mM 2DG/10 μM CCCP for 20 min and subsequently treated with 2 μM BMH-21 for 5 min in the presence of 2DG/CCCP.

### Etoposide is less effective for ATP-depleted cells

Finally, we examined whether 2DG/CCCP treatment could affect the cytotoxicity of etoposide in comparison with two additional DNA-damaging agents, mitoxantrone (MTX) and neocarzinostatin (NCS). DNA damage induced by etoposide is thought to be totally dependent on the presence of functional Top2 ^43^. MTX and NCS act as DNA-damaging agents in a Top2-independent manner ^44,45^. As shown in Fig. 8, etoposide treatment without 2DG/CCCP caused significant reductions in cell viability. The same concentrations of etoposide in the presence of 2DG/CCCP were less effective, and we observed higher cell viability compared to that without 2DG/CCCP. MTX and NCS exhibited similar cytotoxicity regardless of the 2DG/CCCP treatment (Fig 8). Collectively, these observations indicate the relationship between cellular ATP status and the potency of etoposide, suggesting therapeutic implications of the nucleolar translocation of Top2.

**Fig. 8.**
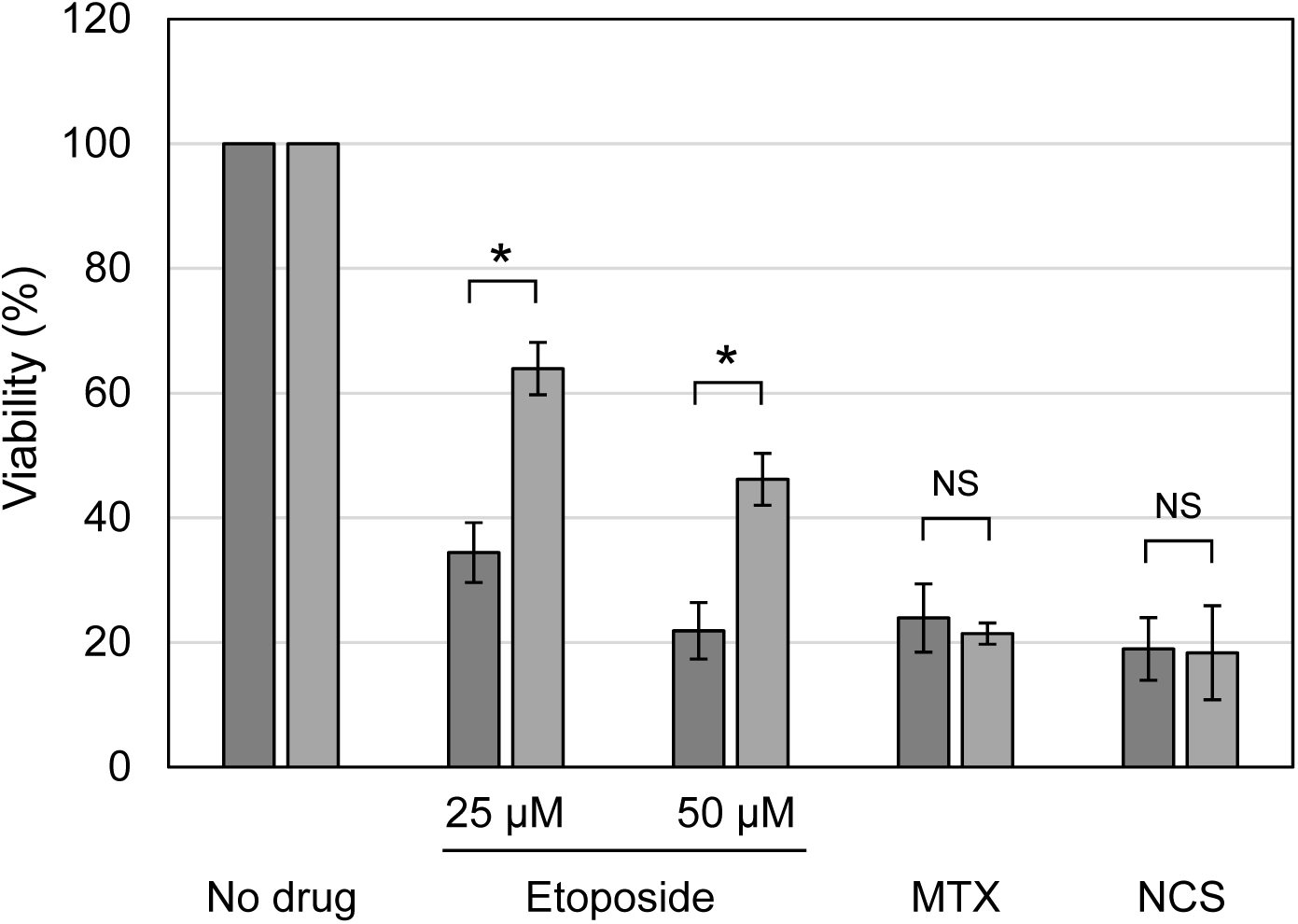
Etoposide is less effective in ATP-depleted cells. HeLa cells were treated with either 25 μM etoposide, 50 μM etoposide, 200 nM mitoxantrone (MTX), or 250 μg/ml neocarzinostatin (NCS) in the presence and absence of 25 mM 2DG/10 μM CCCP for 30 min. Cells were washed and incubated in medium without drugs for 3 days. Average values of viability and SD were calculated (n=5). Statistical values are as follows; *: statistically significant, 25 μM etoposide p=6.63E-06, 50 μM etoposide p=2.11E-05. NS: Not significant, MTX p=0.375, NCS p=0.882.

## Discussion

Previous studies have demonstrated that ATP depletion augments the nucleolar localization of Top2B ^15,20,23^, which represents an important instance that cellular ATP status affects the balance of protein distribution between the nucleoplasm and nucleolus. To date, however, much less attention has been paid to the molecular details of Top2 behaviors with relevance to ATP status, as compared to other well-characterized examples for stress-induced shifts in the nucleoplasm-nucleolus balance, such as p53 and WRN. In this study, we extensively analyzed the nuclear dynamics of Top2A and Top2B in ATP-depleted cells, which significantly extended the previous findings on Top2B. We observed that both Top2A and Top2B quickly translocate from the nucleoplasm to the nucleolus by ATP depletion. FRAP analysis demonstrated that both proteins are highly mobile in the nucleoplasm and nucleolus. These observations suggest that Top2 exists in a certain dynamic equilibrium between the nucleoplasm and nucleolus and that ATP depletion alters this balance. Remarkably, the retention of Top2 in the nucleolus was extremely sensitive to BMH-21. The subnuclear behaviors of the two proteins are essentially indistinguishable, presumably owing to their similarities in domain structure and amino acid sequence, although they are differentially expressed and are involved in different nuclear functions.

This study raises several important questions that provide potential directions for further investigation. First, the interaction partner(s) of Top2 in the nucleolus should be identified in future studies. The retention of mobile proteins in the nucleolus is generally considered to involve the interaction with less mobile nucleolar components, including rDNA, its transcripts, and proteins ^46^. Based on the shift of Top2 distribution from the nucleoplasm to the nucleolus by ATP depletion, we speculate that low ATP conditions may lead to increased interactions of Top2 with some nucleolar components. Our observations on the potent inhibitory effect of BMH-21 suggest that the target of BMH-21 is the interaction partner of Top2A and Top2B in the nucleolus. BMH-21 is a Pol I inhibitor that exerts its inhibitory effect by intercalating into DNA GC-rich regions, particularly G-quadruplex structures, in rDNA ^47,48^. Thus, we speculate that Top2 proteins associate with either some component(s) of the Pol I machinery or GC-rich regions of rDNA. Future research should aim to identify a nucleolar component that shows increased association with Top2 under low ATP conditions.

The second question is how ATP status affects the equilibrium of Top2 distribution between the nucleoplasm and nucleolus. Previous studies have demonstrated that several protein modifications are induced under stressed conditions and contribute to alterations in subnuclear protein distribution. For example, PI3-kinase is activated by proteasome inhibition and facilitates the nucleolar translocation of MDM2 and p53 ^28^. The nucleoplasm-nucleolus distribution of WRN is controlled by catalytic activities of ATM, HDAC, and Sirt1 ^35,36^. Previous studies reported various post-translational modifications of Top2 ^49-51^, some of which may be induced by low ATP, leading to increased interactions between Top2 and nucleolar components. Another, yet not mutually exclusive, possibility is that low ATP status alters the material properties of Top2 and/or its interacting partners in living cells. Although micromolar concentrations of ATP are sufficient for most ATP-driven catalytic reactions, human cells maintain much higher ATP levels on the order of millimolars ^52^. A recent study revealed that physiological concentrations of ATP serve as a natural hydrotope that increases protein solubility in aqueous phases and counteracts the phase separation of macromolecules ^52^. We infer that low ATP levels may alter the properties of Top2 and/or its interacting partners in the cellular milieu, leading to enhanced interactions of them and increased localization of Top2 in the nucleolus. In this respect, it would be interesting to examine the effect of different concentrations of ATP on the material properties of Top2 in the aqueous phase.

The third question is the biological meaning of the nucleolar translocation of Top2 by ATP depletion. We speculate that the translocation of Top2 by ATP depletion may reflect a certain part of the nuclear proteostatic mechanism known as nucleolar sequestration ^37-39^. The microenvironment in the nucleolus is thought to be protective against irreversible aggregation and proteolysis ^37,53^. Nucleolar sequestration shields a fraction of nuclear proteins from the adverse effects of environmental and metabolic stress and serves as a fundamental mechanism for protein homeostasis in the nucleus ^37-39^. Disturbance of proteostasis in the nucleolus was reported to cause the formation of toxic aggregates, which is implicated in the pathogenesis of neurodegenerative disorders ^37,39,54^. In addition, we assume that Top2 may be prevented from accessing its sites of action under ATP-depleted conditions. Top2 has an ATPase domain, and its activity is driven by ATP hydrolysis. Because Top2 is involved in various nuclear processes, the lack of sufficient ATP may cause unfavorable actions of Top2 in some nuclear contexts. Thus, the transient sequestration of Top2 from its primary sites of action in the nucleoplasm may be beneficial during the restoration of ATP levels.

Finally, it is worth noting the intriguing similarities between the nucleolar Top2 and biomolecular condensates formed by liquid-liquid phase separation (LLPS). LLPS drives the compartmentalization of a specific set of biological molecules in a condensed phase, leading to the formation of a biomolecular condensate in a confined space ^55,56^. Molecules in the condensate are generally mobile and exist in a dynamic equilibrium with the surrounding milieu ^55,56^. LLPS-derived condensates are often highly sensitive to the alcohol 1,6-hexanediol and are instantly dissolved by this compound ^57,58^. ATP serves as a hydrotope and counteracts the action of LLPS ^52^. As for the Top2 proteins, they are highly mobile and exist in a certain equilibrium between the nucleolus and nucleoplasm. ATP depletion shifts this balance to the nucleolus side. Addition of BMH-21 rapidly disrupted the retention of Top2 in the nucleolus. Notably, the nucleolus consists of multilayered condensates that possess characteristics of LLPS ^59,60^. Further research with emphasis on the similarities and differences to LLPS will shed more light on the Top2 dynamics in living cells.

## Materials and Methods

### Reagents

2-deoxy-glucose, α-amanitin, etoposide, and mitoxantrone were obtained from FUJIFILM Wako Pure Chemical (Japan). Carbonyl cyanide m-chlorophenylhydrazone and neocarzinostatin were purchased from Sigma-Aldrich (USA). ML-60218 was obtained from Focus Biomolecules (USA). BMH-21 was obtained from Cayman Chemical (USA). BMH-21 was dissolved in DMSO at 2 mM, dispensed into small aliquots, and stored at -80C until use. Once thawed, an aliquot of BMH-21 was not frozen again. BMH-21 was kept away from light during preparation and experiments.

### Plasmids

EGFP-Top2A expression plasmid was constructed in this study. Briefly, total RNA was extracted from HCT-116 cells by using an RNAiso plus reagent (Takara Bio, Japan) and was subsequently subjected to reverse transcription followed by PCR using a OneStep RT-PCR Kit (QIAGEN) with primers for human Top2A. Full-length Top2A cDNA was cloned into pCD3F2-EGFP plasmid (a gift from Prof. David Chen, University of Texas Southwestern Medical Center at Dallas) using standard DNA cloning methods. Integrity of the obtained plasmid was validated by sequencing, and the nucleotide sequence of the cloned Top2A cDNA was confirmed to be identical to the NCBI Reference Sequence NM_001067.4 (human Top2A). Primer sequences used for RT-PCR and detailed procedures of the plasmid construction are available upon request. The expression plasmid for EGFP-tagged human Top2B was described previously ^15^. Expression plasmids for EGFP-Nucleolin and EGFP-Fibrillarin were obtained from Addgene (#28176 and #26673, respectively).

### Cell culture and transfection

HeLa cells were obtained from RIKEN BioResource Research Center (Japan). Cells were grown in αMEM supplemented with 10% fetal bovine serum (FBS), 100 μg/mL streptomycin, and 100 units/mL penicillin at 37°C under a humidified condition with 5% CO_2_. For plasmid transfection, cells (6 × 10^4^) were plated in a 35 mm glass-bottomed dish (Matsunami, Japan). After 24 hr incubation at 37°C, a plasmid (300 ng) was transfected using a FuGENE6 reagent (Promega, USA) according to the manufacturer’s instructions. At 48 hr after transfection, cells were used for live cell imaging and FRAP analysis.

### ATP measurement

Cells were plated in a 96-well plate (5 × 10^3^ cells/well). After 24 hr incubation at 37°C, medium was removed, and fresh alphaMEM containing 25 mM 2DG and 10 μM CCCP was added to each well. Cell lysis and ATP measurement were performed using a CellTiter-Glo luminescent cell viability assay kit (Promega) according to the manufacturer’s instructions. Luminescence was measured using a 2030 ARVO X multilabel reader (Perkin Elmer, USA).

### Immunofluorescence staining

Cells were fixed with 4% paraformaldehyde dissolved in Dulbecco’s phosphate-buffered saline (D-PBS) for 15 min at 4°C. After washing with D-PBS, cells were permeabilized with 0.4% Triton X-100 in D-PBS at room temperature for 5 min. Cells were subsequently blocked with 1% bovine serum albumin in D-PBS for 15 min and reacted with appropriate primary antibodies overnight. After washing with D-PBS, cells were incubated with fluorescent secondary antibodies and subsequently mounted in a Vectashield mounting medium with DAPI (Vector Laboratories, USA). Antibodies used in this study were as follows: mouse anti-Top2A monoclonal antibody (M042-3S, Medical and Biological Laboratories, Japan), mouse anti-Top2B monoclonal antibody (611492, BD Biosciences, USA), rabbit anti-Fibrillarin monoclonal antibody (2639S, Cell Signaling Technology, USA), goat anti-mouse IgG-Alexa Fluor488 antibody (A11029, Thermo Fisher Scientific, USA), goat anti-rabbit IgG-Alexa Fluor594 antibody (A11037, Thermo Fisher Scientific).

### Fluorescence microscopy

Fluorescence microscopy was performed as described previously ^15^. Briefly, an FV1200-IX83 laser scanning confocal microscope with an oil-immersed 60× objective (Olympus, Japan) was used for fluorescence microscopy. For live cell imaging, cells grown on a glass-bottomed dish were placed on a stage top incubator (Tokai Hit, Japan) that maintained a humidified atmosphere of 5% CO_2_ at 37°C. Images were captured and analyzed using FLUOVIEW software (Olympus). Fluorescence intensity of a region of interest (ROI) was quantified using FLUOVIEW software.

### FRAP analysis

FRAP analysis was carried out as reported previously ^15^. In brief, live cell imaging and fluorescence quantification were performed using an FV1200-IX83 microscope and FLUOVIEW software as described above. A spot in either the nucleoplasm or the nucleolus was photobleached with a 473 nm laser at 35% output for 1 sec. Before and after photobleaching, fluorescence images were acquired at 5 sec intervals. All measurements in the photobleached areas were corrected for nonspecific bleaching during monitoring with reference to an unbleached area in the same cell. Fluorescence intensities before and after photobleaching were set to 1.0 and 0, respectively, and fluorescence recovery was expressed as a ratio of fluorescence before and after photobleaching.

### Cell viability measurement

Cells were plated in a 96-well plate (1,000 cells/well) and incubated at 37°C for 24 hr. Thereafter, cells were treated with an appropriate DNA damaging agent with or without 25 mM 2DG/10 μM CCCP at 37°C for 30 min. Subsequently, drug-containing medium was removed, and cells were washed with medium. Incubation was continued in fresh medium at 37°C for 72 hr. Cell viability was measured using a Cell Counting Kit-8 (Dojindo, Japan).

### Statistical analysis

Statistical analysis was performed using Welch’s t-test. The number of experiments and p-values are indicated in the figure legends.

## Data availability

The data that support the findings of this study are available from the corresponding authors upon reasonable request.

## Acknowledgements

This work was supported by JSPS KAKENHI Grant Number 19H04271 to K.i.Y.

## Author contributions

K.M.Y. and K.i.Y. conceived the study, designed and performed experiments. K.M.Y. and K.i.Y. drafted, reviewed, and approved the manuscript.

## Competing interests

The authors declare no competing interests.

## Notes

### Competing Interest Statement

The authors have declared no competing interest.

